# Identification of a novel efficient transcriptional activation domain from Chinese fir (*Cunninghamia lanceolata*)

**DOI:** 10.1101/2020.04.20.051466

**Authors:** Tengfei Zhu, Wenyu Tang, Delan Chen, Renhua Zheng, Jian Li, Jun Su

**Affiliations:** Basic Forestry and Proteomics Research Center, College of Forestry, Fujian Provincial Key Laboratory of Haixia Applied Plant Systems Biology, Fujian Agriculture and Forestry University, Fuzhou 350002, China; Bureau of Forestry, Wuyishan, Fujian 354300, China; Fujian Academy of Forestry, the Key Laboratory of Timber Forest Breeding and Cultivation for Mountainous Areas in Southern China, State Forestry Administration Engineering Research Center of Chinese Fir, the Key Laboratory of Forest Culture and Forest Product Processing Utilization of Fujian Province, Fuzhou 350012, China; College of Forestry, Fujian Agriculture and Forestry University, Fuzhou 350002, China

**Keywords:** Transcriptional activation domain, Chinese fir (*Cunninghamia lanceolata*), TAC3, VP16, EDLL, SRDX, Flowering Locus C (FLC), Flowering Locus T (FT), Peptide transformation

## Abstract

Activation domains are used as critical components of artificial gene modification tools for genetic breeding. The high efficiency of the activation domain relies on the host plant. However, no activation domain has been identified that originates from Chinese fir (*Cunninghamia lanceolate*). In this study, a novel strong activator was identified from the whole Chinese fir cDNA library. This plant conserved activator was named TAC 3 (Transcriptional Activation domain from Chinese fir 3). C-terminal 70 amino acids of TAC (TAC3d) have a stronger ability than the commonly used strong activation domain of the virus protein VP16, or the strong plant activation domain, EDLL, in Chinese fir. Through Dual-luciferase assay, phenomic analysis and FT (Flowering Locus T [FT]) quantification, it was shown that, TAC3d can overcome the transcriptional repression of strong plant repressors (Flowering Locus C [FLC]) when fused to its C-terminal domain, thus inhibit the repression of FT expression. In conclusion, for the first time, an activation domain has been identified from Chinese fir. TAC3, which can be used for precise gene activation in Chinese fir in the future, and its function in the plant is more powerful than the commonly used strong activation domain (such as VP16 and EDLL).

**Highlight:** TAC3 is the first transcriptional activation domain identified from Chinese fir and its function is more powerful than some commonly used strong transcriptional activators (such as VP16 and EDLL)

## Introduction

Direct manipulation of gene expression in vivo is a powerful approach for investigating biological systems and bioengineering in plants (Yueying *et al*. 2015, Levi *et al*. 2018). In the last decade, considerable progress has been made in terms of the development of techniques for gene targeting. Synthetic transcriptional activators have been generated by fusing programmable DNA-binding modules to a variety of activation domains. Such DNA-binding modules include zinc-finger domains, transcriptional activator-like (TAL) effectors, and clustered regularly interspaced short palindromic repeats (CRISPRs) and CRISPR-associated (Cas) regulatory systems. These synthetic transcriptional activators are used to selectively activate target genes to improve multiple plant traits (Bartsevich *et al*. 2003, Li *et al*. 2013, Agnieszka *et al*. 2015).

The upregulation of gene expression level by synthetic transcriptional activators relies on the activation ability of their own activation domains. The strong activation domain that is commonly used in plants is the herpes simplex virus early transcriptional activator, VP16, and its four tandem repeats VP64 (Scott *et al*. 2014). However, several activation domains that originate from plants have also been identified and well characterized, such as the EDLL motif (Tiwari *et al*. 2012) and the TAL activation domain (Mahfouz *et al*. 2011). In contrast to activation domains from plants, VP16 has not always worked efficiently in plants, especially when fused to plant repression motifs (Ohta *et al*. 2001, Sumire *et al*. 2014). This is mainly because VP16 was originally isolated from a human cell, which has a different working environment to a plant cell (Blaise *et al*. 2016). Further identification of stronger plant-originated activation domains is crucial for the precise activation of gene expression in target plants.

Chinese fir (*Cunninghamia lanceolata* [Lamb.] Hook) is the second most dominant plantation tree species for afforestation in the world. It is widely planted in southern China and composes approximately 21.35% of Chinese plantations (Yuan *et al*. 2009, Xian *et al*. 2017). Although the rotation period of Chinese fir plantations has been shortened to 20–25 years, it is still difficult to solve obstacles such as acidic soil and drought and pest damage through traditional Chinese fir breeding methods (Tian *et al*. 2011). Many candidate genes related to the stress and quality of Chinese fir timber have now been identified, and a regeneration system has been established for Chinese fir. Therefore, it is now possible to improve Chinese fir traits via gene modification, which will greatly shorten the breeding period (Xu *et al*. 2019, Zhihui and Sizu 2019). However, a strong activation domain that originates from Chinese fir, which is crucial for the genetic modification of Chinese fir, is yet to be found.

In this study, a strong activation domain was identified from a Chinese fir cDNA library. The transcriptional activation ability of the domain was verified using both in vivo and in vitro assays of Chinese fir. Furthermore, the possible applications of this activation domain were tested. To the best of our knowledge, this study is the first to identify an activation domain from Chinese fir and to characterize the domain in vivo. This work provides the final component for a gene modification tool that can be used in Chinese fir.

## Materials and methods

### Chinese fir cDNA library preparation

The procedure of generating a full-length cDNA library has previously been described (Kooiker and Xu 2014); the library was constructed using 1 μg of equally mixed RNAs. First-strand cDNA was synthesized using a Clontech SMARTer PCR cDNA Synthesis Kit (cat. no. 634926, www.clontech.com/) with anchored oligo(dT)30 primer. Double-strand cDNA was amplified by long-distance PCR (LD-PCR) using an Advantage 2 PCR kit (Clontech, cat. no. 639206).

### Reporter and effector constructs

The reporter constructs are described in the figures. The reporters were inserted into pGreenII0800 backbones using a BamHI restriction enzyme digestion site. The promoter sequence of *Flowering Locus T* (*FT*) has been described previously (Huang *et al*. 2005), as has the sequence of the Gal4 upstream activating sequence (Gal4UAS) (Tiwari *et al*. 2012).

All effector genes contained a double CaMV 35S promoter followed by a translational enhancer and a nopaline synthase (NOS) terminator. These genes were constructed into pEarleygate104 using a gateway system. All the sequences of the effector genes are listed in Table S1. All effector genes constructed with the yeast Gal4 DNA-binding domain (amino acids [aa] 1–147) also contained an N-terminal fusion of four copies of the c-Myc epitope.

### Protoplast transformation assay

Isolation and infection of mesophyll protoplasts from Arabidopsis (*Arabidopsis thaliana*) leaves were performed as described previously (Tiwari *et al*. 2012). The co-transfection assays were performed using 15 µg of reporter plasmid and 10 µg of effector plasmid. All transfections were conducted in triplicate, and three independent transfections were performed for each set of experiments.

### Dual-luciferase assay

The dual-luciferase assay was conducted by transferring the relative constructs into mesophyll protoplasts from Arabidopsis leaves. The corresponding Firefly luciferase (LUC) and Renilla luciferase (REN) values were measured 16 h after transfection using a Dual-Luciferase® Reporter Assay System (Promega, WI, USA).

### Developing the transgenic plants, flowering time analysis, and gene expression quantification

Transgenic lines were prepared that overexpressed yellow fluorescent protein-flowering locus C (YFP-FLC) or YFP-FLC-activation domain (AD). To prepare these lines, the coding regions of *AtFLC* with or without C-terminal fusion of the AD were cloned into a pEarleygate104 vector using a gateway system to generate the related construct. These constructs were then transferred into *Agrobacterium tumefaciens* (strain AGL0), which was allowed to infect Col-0 Arabidopsis using the standard floral-dip method. The transgenic T1 populations were screened on compound soil sub-irrigated with the BASTA solution.

The flowering time was determined as the number of days from planting to the appearance of the first visible bud (of the T2 plant). The numbers of rosette leaves were counted at the showing of the first visible bud.

Total RNAs were isolated from Arabidopsis plants using Trizol reagent (Invitrogen, USA). Total RNA (1 mg) was reverse transcribed using the PrimeScript RT Reagent Kit with the gDNA Eraser (TAKARA, Japan). Quantitative PCR reactions were performed using the gene-specific primers of *FLC* and *FT* (Table S1) with Hieff™ qPCR SYBR® Green Master Mix (Low Rox Plus, China) on a QuantStudio 6 Flex PCR (Life Technologies Corporation, CA, USA). The qPCR signals were normalized to that of the reference gene *PP2A* in Arabidopsis by using the ΔCT method. Biological triplicates with technical duplicates were used in all cases.

## Results

### Identification of Chinese fir transcriptional activation domain (TAD)

A TUP1 conjugated activation domain (AD) screening system was used for identifying the TAD from the Chinese fir full-length cDNA library (approximately 1 × 10^5^ genes); 3 million single colonies were screened (Fig. S1). Only seven single colonies survived, and they contained Transcriptional Activation domain from Chinese fir 1 (TAC1), TAC2, and TAC3 (Table S1). Both TAC1 and TAC3 were characterized as TADs in yeast and displayed stronger activities than VP16 (Fig. S2). However, only TAC3 overcame the strong suppression of the yeast general transcriptional corepressor TUP1 (Fig. S3). Furthermore, by measuring the activation activity of its short fragments in yeast, it was elucidated that the C-terminal end of the TAD of TAC3 (110–179 aa) had the highest activation activity (Fig. S4).

To test whether TAC3 acted as a TAD in Chinese fir, a peptide transformation assay (Lakshmanan *et al*. 2013) was performed for transient expression of the related constructs in Chinese fir seedlings (Fig. S5). TAC3 was found to be a TAD with higher activity to VP16 (Fig. 1B). However, the co-transfection efficiency was too low in the Chinese fir seedlings, and thus the Arabidopsis transient expression system was adopted for further study.

**Fig. 1.**
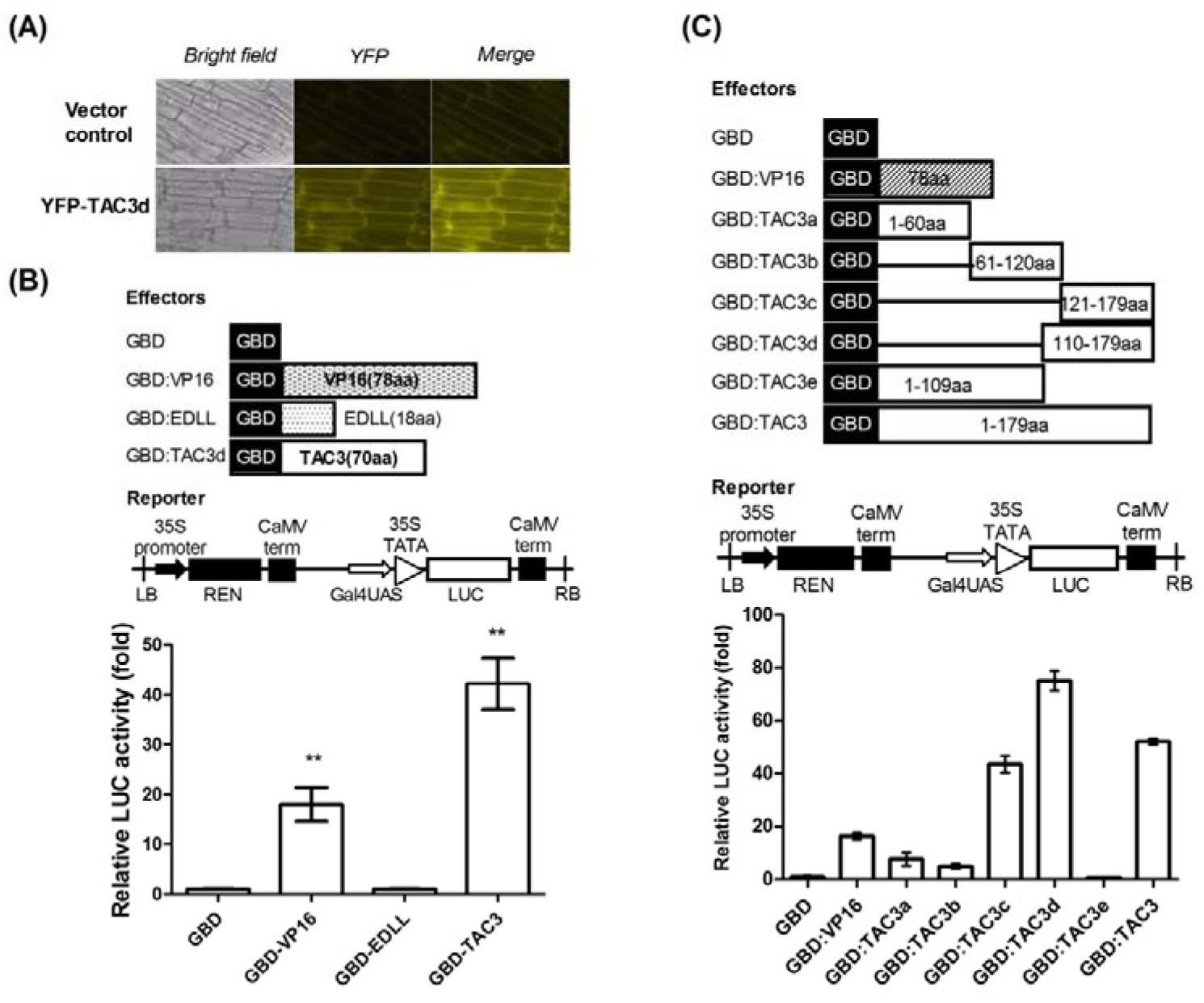
The C-terminal region (TAC3d) of TAC3 is sufficient for stronger transcriptional activity than VP16 in Chinese Fir. (**A**) Expression pattern of TAC3d in Chinese fir. TAC3d was overexpressed in Chinese fir seedlings with N-terminal fusion of yellow fluorescent protein (YFP) using a modified transient expression assay (Fig. S5). (**B**) Different activation domains (TAC3/EDLL/VP16) were separately fused to the C-terminal end of Gal4 DNA-binding domains (GBDs) and co-transferred into Chinese fir seedlings with Gal4UAS::LUC as the reporter. Ten-day-old Chinese fir seedlings were incubated in transfection media with a ratio of peptide/plasmid (w/w) of 0.5, at 25°C and 0.1 MPa for 5 min. Further incubation was conducted in the dark for 2 days. Data are the mean values of firefly luciferase (LUC) activity divided by Renilla luciferase (REN) activity ± SE of nine replicates. ** represents a significant difference at *P* < 0.01 and was detected using one-way ANOVA tests between the samples and negative control. (**C**) The 110–179 amino acid (aa) fragment of TAC3 shows higher transcriptional activity than full-length TAC3. Different fragments of TAC3 were constructed into effectors, with the aim of pinpointing the activation domain. Both effectors and reporters were co-transfected into ten-day-old Chinese fir seedlings. The 35S promoted *REN* gene encoding the Renilla Luciferase served as a non-specific reporter gene. Data are the mean values of Firefly luciferase (LUC) activity divided by Renilla luciferase (REN) activity ± SE of nine replicates.

### The C-terminal region of TAC3 is sufficient for stronger transcriptional activity than VP16

Three Chinese fir TADs (TAC1, TAC2, and TAC3) were individually fused to the C-terminus of Gal4 DNA-binding domains (GBDs). These constructs were then co-transferred with the reporter LUC, which was driven by Gal4UAS, into Arabidopsis protoplasts. The activation ability of the three TADs was then assessed in vivo using a dual-luciferase assay. GBD alone was used as a negative control, and a fusion of GBD with the universal activation domain VP16 was used as a positive control. The full-length TAC3 protein activated the transcription of the reporter by three times the amount that the VP16 activation domain did in planta (Fig. 1A and B).

Fragmentation of the full-length TAC3 sequence into TAC3a (1–60 aa), TAC3b (61–120 aa), and TAC3e (1–109 aa) resulted in total loss of transcriptional activity. However, TAC3c (121–179 aa) displayed a similar transcriptional activation activity to TAC3. This suggests that TAC3c is solely responsible for the activation activity of the TAC3 protein. Moreover, the activation activity of TAC3d (110–179 aa) was 1.5 times greater than that of the full-length TAC3. TAC3d was able to activate the transcription of the reporter to a level that was more than four times that activated by the very strong VP16 activation domain.

### TAC3d is an acidic-type activation domain and can activate transcription from both proximal and distal positions

Transcriptional activation domains can be assigned into three major categories, acidic, glutamine-rich, and proline-rich, which are defined according to the frequency of specific amino acids (Kunzler *et al*. 1994, Remacle *et al*. 1997). The 70 aa of the C-terminal TAC3d has an arrangement of acidic amino acids that is unusual and differs from other well-defined acidic activation domains such as Gal4 and VP16 (Triezenberg *et al*. 1988, Blair *et al*. 1994). It has been shown that for the plant acidic activation domain, EDLL, its net level of gene activation not only depends on the potency of the transcriptional activation domain but also on the concentration of the activator and the number of DNA-binding sites (Tiwari *et al*. 2012).

The activation potency of TAC3d was tested for using heterologous LUC reporters containing either single or multiple copies (4X) of the Gal4UAS. EDLL motifs and VP16 were used as positive controls. As shown in Fig. 2A, VP16, the EDLL motif, and TAC3d were all able to activate the transcription of the single Gal4UAS::LUC reporter gene to different extents above GBD alone: VP16, 47-fold; EDLL, 9-fold; and TAC3, 90-fold. Furthermore, the activity of the EDLL motif (39-fold) was slightly lower than that of VP16 (46-fold) when multiple DNA-binding sites were used (Fig. 2B). Meanwhile, TAC3d (149-fold) displayed more than three times the activity of VP16 when multiple DNA-binding sites were used. There was no significant difference in the activity of the VP16 activation domain between the assays using single or multiple DNA-binding sites (Fig. 2A and B). However, obvious changes were recorded for EDLL and TAC3d.

**Fig. 2.**
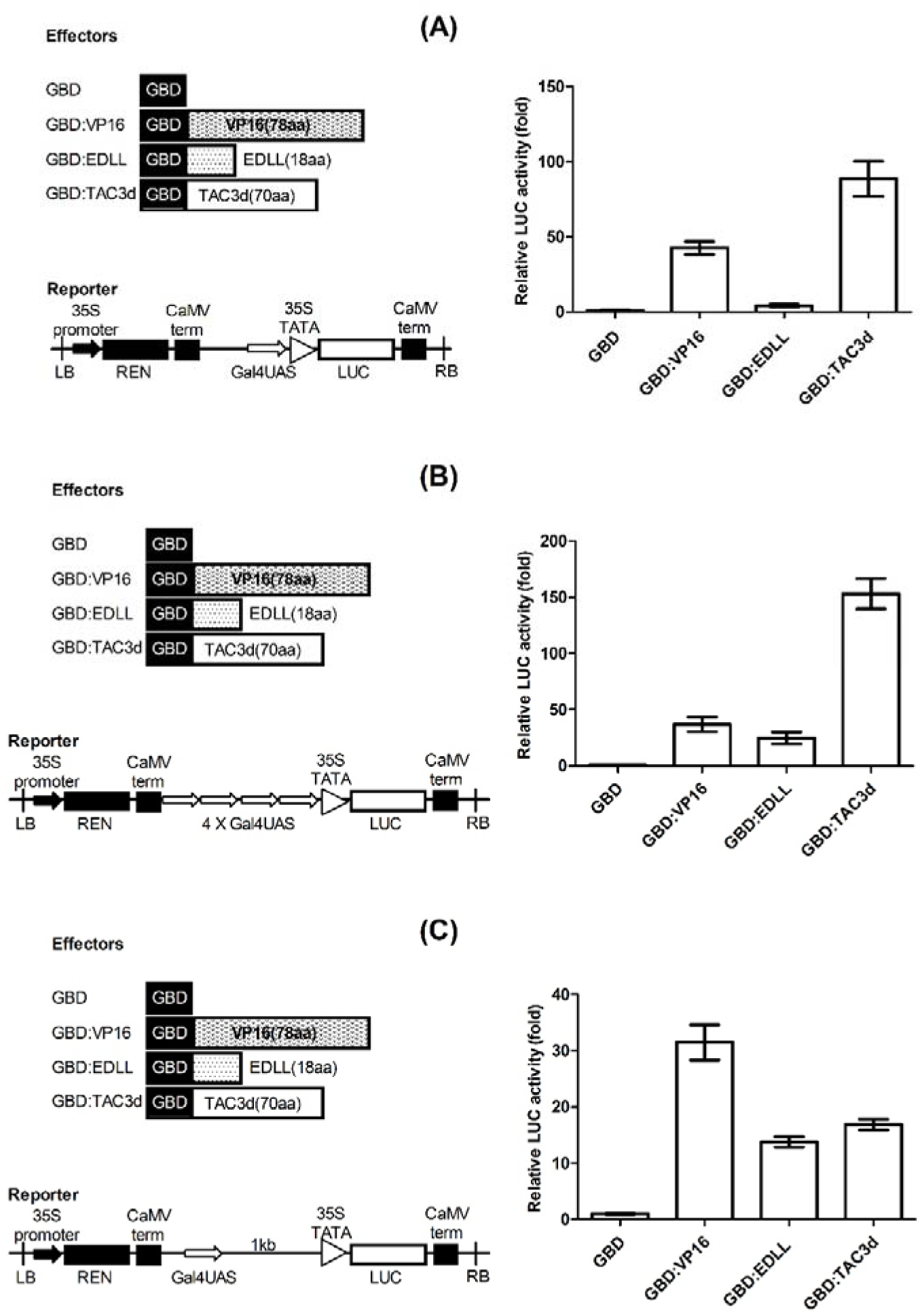
TAC3 can activate transcription from both TATA-proximal and remote positions. Co-transfection was performed in Arabidopsis mesophyll protoplasts using different effectors and reporters. Firefly luciferase/Renilla luciferase (LUC/REN) represents the activation ability relative to the Gal4 DNA-binding receptor (GBD) effector. For each effector, three repeats were measured, and the value is the mean ± SD. (**A**) Using *Gal4UAS::LUC* as the reporter gene, a single copy of Gal4 DNA-binding sequence (Gal4UAS) was inserted into the reporter in a position that was proximal to the 35S minimal TATA box. (**B**) Using *4XGal4UAS::LUC* as the reporter gene, quadruple Gal4UAS was inserted into the reporter in a position that was proximal to the 35S minimal TATA box. (**C**) Using *Gal4UAS1Kb::LUC* as the reporter gene, a single copy of Gal4UAS was inserted into the reporter 1 kb upstream of the 35S minimal TATA box.

Several studies have shown that activation domains differ significantly in their ability to drive transcription according to their binding position relative to the TATA box in the promoter region. Acidic activation domains (including EDLL) are functional at both proximal and distal positions (Seipel *et al*. 1992, Kunzler *et al*. 1994, Remacle *et al*. 1997, Tiwari *et al*. 2012). To determine whether TAC3 has a similar function to other acidic activation domains, an artificial reporter was created, in which a single Gal4UAS binding site was present 1 kb upstream of the 35S TATA box (Fig. 2C). Interestingly, VP16 was almost equally active from both proximal and distal positions within the promoter. However, the activities of both EDLL and TAC3d were significantly reduced at the distal position (EDLL, 11-fold and TAC3d, 15-fold), but transcription was still promoted (Fig. 2C).

### The activation function of TAC3 is conserved across plant species

A phylogenetic analysis was carried out for TAC3 from Chinese fir (ClTAC3) and its orthologs from other plant species (Fig. 3A). It was found that ClTAC3 was more closely related to the TAC3 of crops (such as rice, maize, and soybean) than of trees. A BLAST search for the C-terminal 70 aa (TAC3d) and full-length TAC3 was performed against the collection of plant genomes at the National Center for Biotechnology Information (http://www.ncbi.nlm.nih.gov/). The alignment of selected orthologs (Fig. S6) highlighted a strongly conserved motif (LLxASDDxLGxP) that was located at 81–92 aa of TAC3, which is a region that has limited activation ability. Meanwhile, TAC3d only had hits with eight unknown proteins from *Picea stenoptera*.

**Fig. 3.**
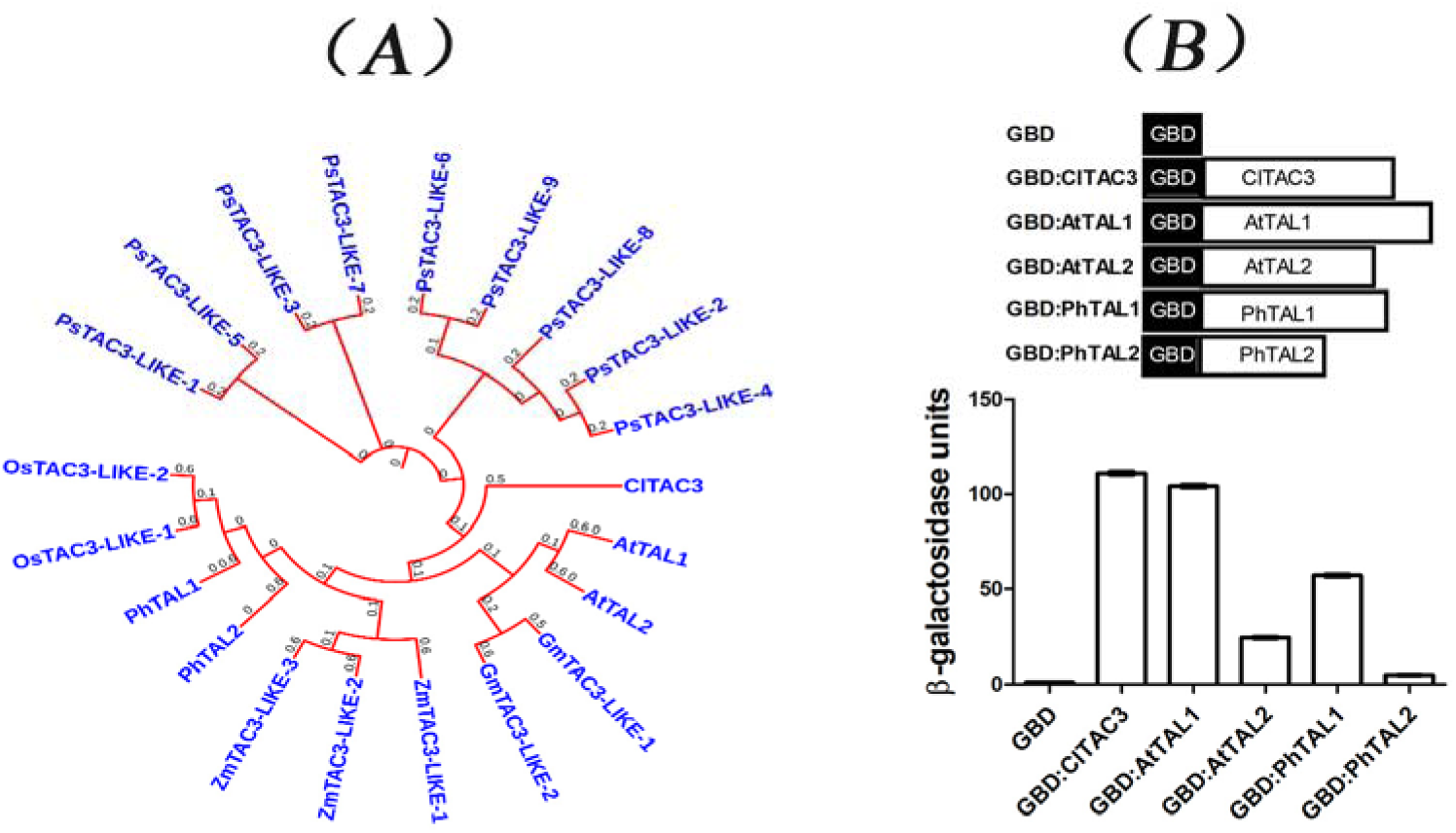
Transcriptional activity of TAC3 and orthologs from other species. (**A**) A phylogenetic tree was built based on the amino acid sequences of TAC3 and its orthologs from *Cunninghamia lanceolata* (Cl), *Pterocarya stenoptera* (Ps), *Arabidopsis thaliana* (At), *Oryza sativa* (Os), *Phyllostachys heterocycle* (Ph), *Glycine max* (Gm), and *Zea mays* (Zm). The phylogenetic tree was constructed from the protein sequences obtained from the National Center for Biotechnology Information (NCBI), and the analysis was performed as described previously (Wenjia *et al*. 2017) and modified online in iTOL (https://itol.embl.de/). (**B**) All the constructs were transferred into the yeast strain Y187. A CPRG assay (Yeast Protocols Handbook, Clontech) was conducted to measure the β-galactosidase activity of the different constructs in yeast. There were three samples for each construct and three repeats for each sample. Boxes and whiskers represent the means and standard errors, respectively.

To test whether the TAC3 orthologs had similar activation functions, two TAC3 members from Arabidopsis (AtTAL1 and 2) and Moso bamboo (*Phyllostachys heterocycle*; PhTAL1 and 2) were fused to the C-terminal ends of GBDs, and transcriptional activity was evaluated in yeast (Fig. 3B). GBD alone was used as the negative control. Compared to the negative control, all five TAC3 members from the different plants were shown to be transcription activators. The activity of ClTAC3 was significantly higher than the activity of the TAC3 orthologs from Arabidopsis and Moso bamboo (ClTAC3, 111-fold; AtTAL1, 103-fold; AtTAL2, 22-fold; PhTAL1, 52-fold; PhTAL2, 6-fold). Although the specific reasons for this higher activity were not further explored, it can be concluded that TAC3 members from other plant species generally also possess activation potential.

### TAC3d can relieve transcriptional repression on heterologous DNA-binding proteins

TAC3d is relatively small (70 aa) compared to commonly used activation domains such as VP16 (78 aa) (Triezenberg *et al*. 1988). To illustrate the additional characteristics and the utility of this activator sequence, we first tested the ability of TAC3d to overcome a conventional plant repressor motif SRDX (Ohta *et al*. 2001, Sumire *et al*. 2014). It was found that fusing TAC3d to the C-terminal end of SRDX reversed its transcriptional repression, and that this effect was much stronger with TAC3d than with VP16 and the EDLL motif (Fig. 4A).

**Fig. 4.**
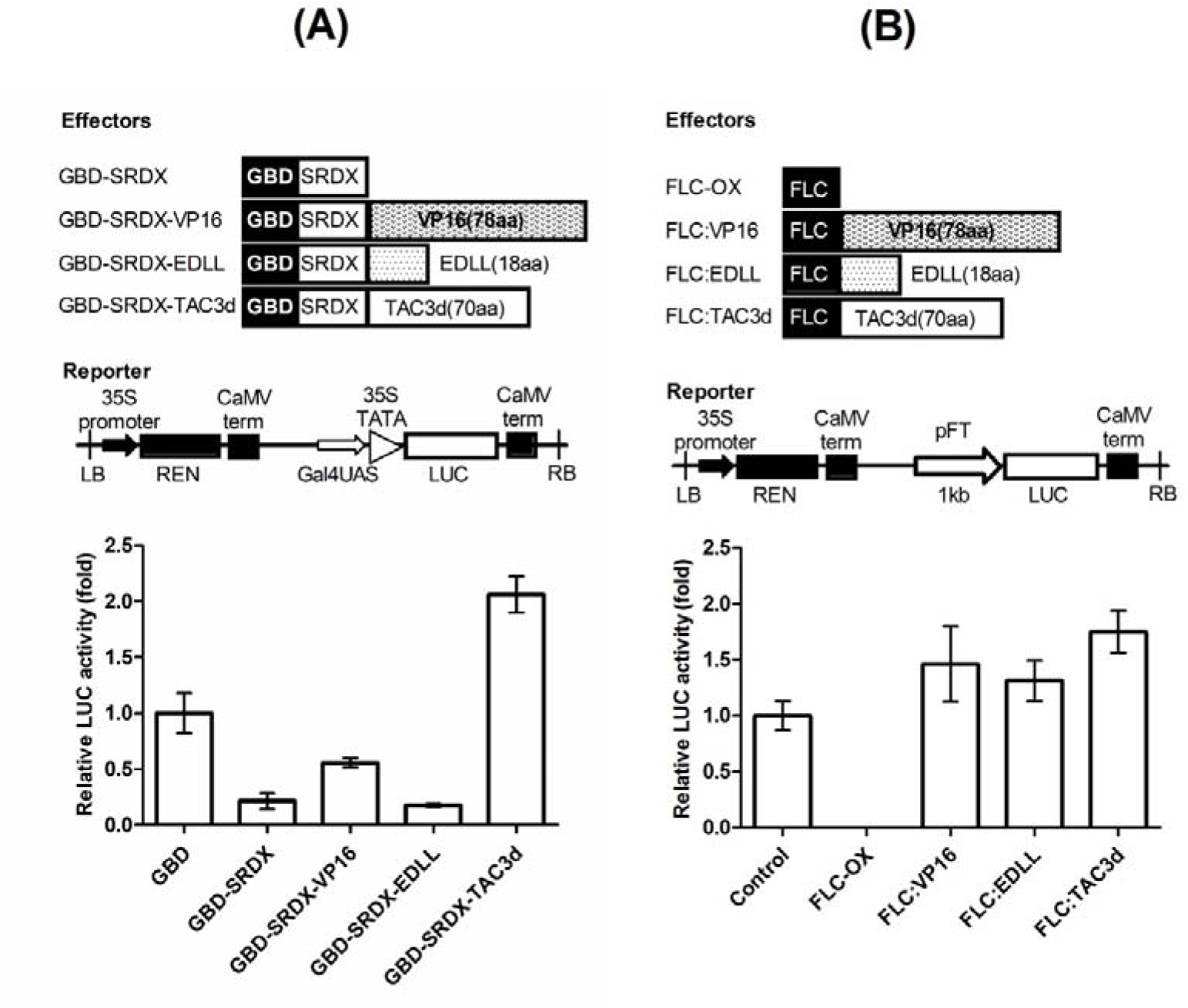
TAC3 can reverse the transcriptional repression caused by repressors. (**A**) Different activation domains (TAC3d/EDLL/VP16) were individually fused to the C-terminus of Gal4 DNA-binding domain (GBD)-SRDX and co-transferred into Arabidopsis mesophyll protoplasts with Gal4UAS::LUC as the reporter. Data are the mean values of Firefly luciferase (LUC) activity divided by the Renilla luciferase (REN) activity ± SD of three replicates. (**B**) Different activation domains (TAC3d/EDLL/VP16) were separately fused to the C-terminus of AtFLC and co-transferred into Arabidopsis mesophyll protoplasts with pFT::LUC as the reporter. Fusion of TAC3d relieved the repression caused by AtFLC on the AtFT promoter. Data are the mean values of Firefly luciferase (LUC) activity divided by the Renilla luciferase (REN) activity ± SD of three replicates.

Flowering is a complex trait in plants, and FLC is a crucial floral regulator that integrates multiple pathways to fine-tune flowering time. The *FLC* gene encodes a MADS-box transcription factor, which acts as a strong repressor of floral transition by directly binding to the promoters of several flowering time genes (including FT) and repressing their mRNA transcription (Michaels and Amasino 1999, Sheldon *et al*. 1999, Ratcliffe *et al*. 2003).

Therefore, we subsequently examined whether such a short activation domain (TAC3d) could be used to relieve the transcriptional repression of a heterologous DNA-binding protein (FLC) that can directly bind to the promoter region of FT to suppress FT expression. To do this, VP16, the EDLL motif, and TAC3d were each individually fused to the C-terminal end of FLC, which targeted these three activation motifs to the reporter (*pFT*::LUC). TAC3d was found to display a significantly greater ability to reverse the suppression caused by FLC than EDLL or VP16 in Arabidopsis protoplasts (Fig. 4B).

The final aim was to further test whether TAC3d could be used as precise gene activation tool in planta. TAC3d, VP16, and the EDLL motif were fused with the C-terminal end of the Arabidopsis FLC protein (FLC-TAC3, FLC-VP16, and FLC-EDLL). These constructs were transferred into Arabidopsis to generate stable transgenic lines, while an FLC overexpression line (FLC-OX) was used for the negative control. FLC-OX and two independent lines of FLC-TAC3, FLC-VP16, and FLC-EDLL were used for further investigation (Fig. 5A). The flowering time and rosette leaf number were quantified from different genotypes. Under long-day conditions, the wildtype (WT) flowered at approximately the 30^th^ day after germination. FLC-OX delayed the flowering time by 5 days compared to the WT. Meanwhile, the other genotypes (FLC-TAC3, FLC-VP16, and FLC-EDLL) reduced flowering times to 15–18 d after germination (Fig. 5B). After flowering, the rosette leaf numbers of the different genotypes were counted (Fig. 5C). Only FLC-OX generated two more rosette leaves than WT (11 rosette leaves), while the six other lines (FLC-TAC3, FLC-VP16, and FLC-EDLL) had only 6–8 rosette leaves after flowering.

**Fig. 5.**
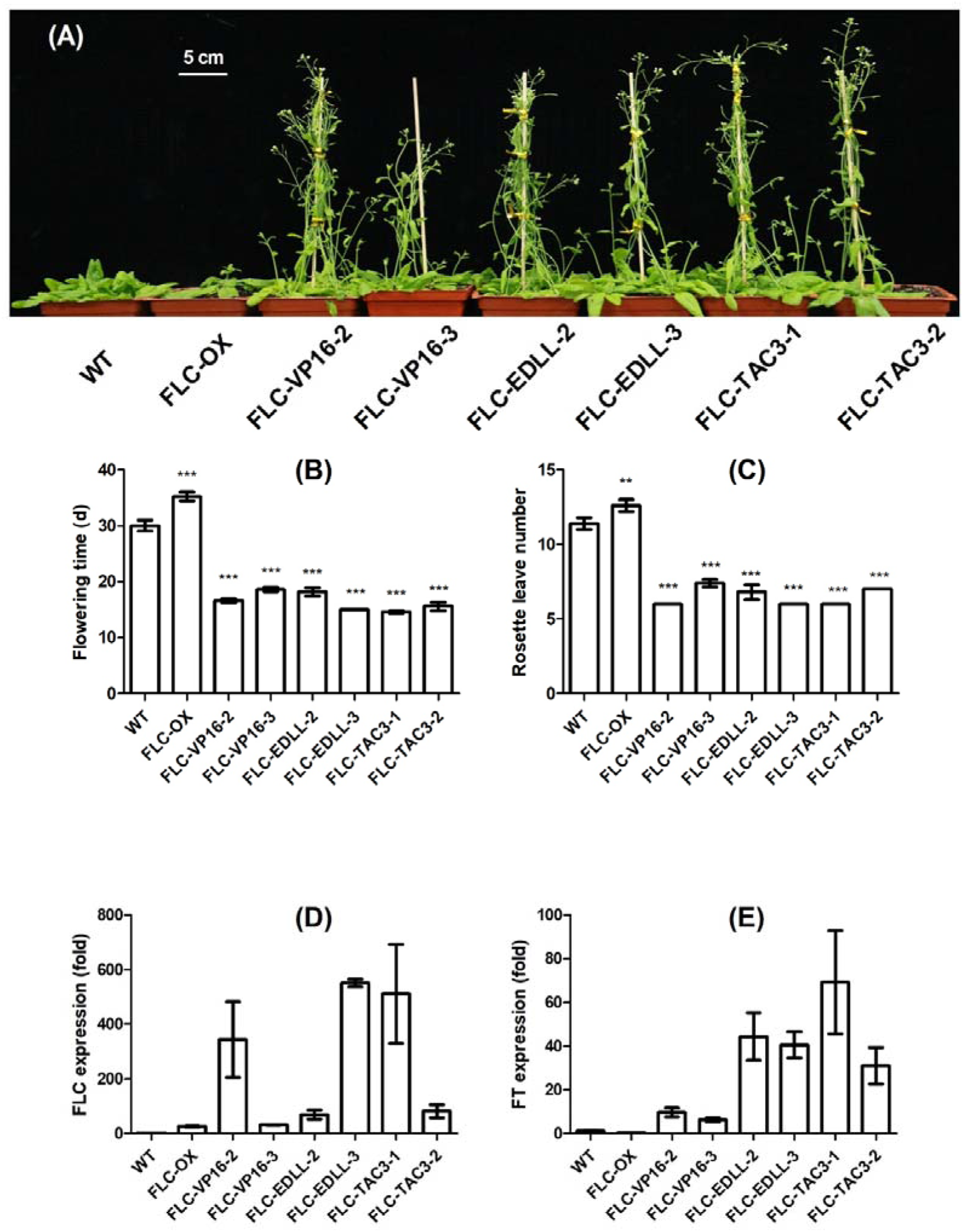
TAC3 fusion rescues the late-flowering phenotype of the FLC overexpressed by upregulating FT expression. TAC3d, VP16, and the EDLL motif were fused to the C-terminus of the Arabidopsis FLC protein with an N-terminal yellow fluorescent protein (YFP) to construct FLC-TAC3, FLC-VP16, and FLC-EDLL, respectively. These were then overexpressed in Arabidopsis. The single FLC overexpression line (FLC-OX) was used as the negative control. (**A**) Three-week-old T2 Arabidopsis transformants of different genotypes (Col-0 background) were grown under long-day conditions (dark:light [D:L] = 8:16). The lines overexpressing FLC-VP16/EDLL/TAC3 flowered earlier than the wildtype (WT). (**B**) Quantification of flowering time and (**C**) rosette leaf number of different genotypes were measured under long-day conditions (D:L = 8:16). Values are shown as mean ± SE of three independent lines, with five replicates for each line. (**D** and **E**) High expression of FLC and FT was detected in lines overexpressing FLC-VP16/EDLL/TAC3. Expression level of (**D**) FLC and (**E**) FT in 10-day-old Arabidopsis seedlings. The expression is shown as the fold change relative to the WT and was determined by quantitative RT-PCR and normalized using *PP2A* expression level. The data are the means ± SD of two independent lines for each genotype. ** and *** indicate statistical differences at *P* < 0.01 and *P* < 0.001, respectively, between samples using one-way ANOVAs and multiple comparison analysis by Turkey’s test.

The expression of FLC and FT in all the independent lines of the different genotypes was also tested (Fig. 5D and E). The expression level of FLC was induced to approximately 24 times that of the WT in the negative control (FLC-OX). However, the FT expression level was strongly suppressed. For FLC-TAC3, FLC-VP16, and FLC-EDLL, both high expression (FLC-VP16-2, 389-fold; FLC-EDLL-3, 564-fold; and FLC-TAC3-1, 544-fold) and low expression (FLC-VP16-3, 20-fold; FLC-EDLL-2, 40-fold; FLC-TAC3-2, 41-fold) lines were chosen according to their FLC expression levels. Both of the independent FLC-VP16 lines showed higher FT expression levels (FLC-VP16-2, 9-fold; FLC-VP16-3, 6-fold) than the WT. This was also true for the FLC-EDLL (FLC-EDLL-2, 44-fold and FLC-EDLL-3, 40-fold) and FLC-TAC3 (FLC-TAC3-1, 69-fold and FLC-TAC3-2, 30-fold) lines. Overall, the above analysis validates the in vivo potency of the C-terminal domain of TAC3 (TAC3d) as a transcriptional activator of a specific gene in planta.

## Discussion

### TAC3d has a stronger activation ability than VP16 conditionally

In this study, the activation efficiency of VP16 was compared in different species (yeast, Arabidopsis, and Chinese fir) using the Gal4-UAS system. This showed that VP16 is a universal activator. Surprisingly, it was found that TAC3 was able to activate target gene expression in both yeast and plants, and its activation ability was much stronger than that of VP16 (Figs. 1 and S2). However, in Arabidopsis, the activation ability of TAC3 was much weaker than that of VP16 in distal positions (Fig. 2). This result was similar to the results found for other plant-specific activation motifs, such as EDLL and TAD (Tiwari *et al*. 2012, Levi *et al*. 2018). These results suggest that activation domains that have originated from plants may not able to work as efficiently away from the TATA box as exogenous activators (such as plant virus activators) (Douglas and Wendy 2015). Therefore, future work must focus on expanding the screening targets from plants to other symbiotic microbes (Afsar *et al*. 2015). In addition, the mechanism of transcriptional activation deserves further exploration.

### TAC3 is a highly conserved novel plant-specific co-activator

In this study, the full length of ClTAC3 was revealed using 3’ and 5’ rapid amplification of cDNA ends (RACE; Table S1). ClTAC3 and its orthologs are proteins of unknown function (Fig. 3), and there is no reference that can be used to predict its function. However, this high transcriptional activation ability of ClTAC3 is unprecedented in plants. From the gene structure, it was determined that the N-terminus of ClTAC3 (1–43 aa) was the nuclear localization site. This was also confirmed by the subcellular localization of green fluorescent protein (GFP)-ClTAC3 in *Nicotiana tabacum* (not shown here). However, no DNA-binding site was identified from any of the orthologs. Altogether, the results of the present study suggested that TAC3 was highly likely to be a transcriptional co-activator.

Interestingly, the middle section of the ClTAC3 protein (81–92 aa) was highly conserved across plant species (Fig. S6) and was similar to the plant transcriptional repression domain SRDX (Sheldon *et al*. 1999). However, the C-terminus of the TAC3 orthologs showed a strong transcriptional activation ability (Fig. 3). These results indicate that TAC3 possesses both a repression and an activation function; the higher activation ability of TAC3d alone compared to that of the full-length protein provides clear evidence for this (Fig. S4). The fusion of repression and activation domains within single proteins has been found in both microbes and mammals, and these proteins have been shown to be functional in response to environmental signals (Ratcliffe *et al*. 2003, Bjorn *et al*. 2006, Rybakova *et al*. 2015).

In conclusion, a novel activation domain (TAC3d) from Chinese fir was identified in this study. The activation activity of this domain was found to be much stronger than that of the widely used activation domain VP16 and the EDLL motif that originated from plants. Our data also suggested that TAC3d was a powerful activator of gene expression in planta. Several research questions remain; the function of TAC3 in the plant should be elucidated, and an explanation for why plants need such a strong activation domain should be sought. Furthermore, it will be important to explore how TAC3d activates gene expression. Our future work on ClTAC3 and its orthologs in Arabidopsis will address these questions.

## Supplementary Data

Table S1. List of primers, cis- and trans-elements used in this research;

Fig. S1. Schematic diagram of the activation domain (AD) screening system in yeast;

Fig. S2. TUP1 represses transcriptional activation in yeast when directly fused to the N-terminus of the Gal4 DNA-binding domain (GBD);

Fig. S3. TAC3 is a stronger transcriptional activator than VP16 in both yeast and Arabidopsis;

Fig. S4. The amino acids 110–179 of TAC3 had a higher activation ability than other TAC3 fragments in yeast;

Fig. S5. Impact of incubation time and ingredient ratio (peptide/plasmid) on transfection efficiency of Chinese fir;

Fig. S6. Amino acid sequence alignments of ClTAC3 and orthologs from other species.

## Acknowledgments

This work was supported by the National Natural Science Foundation of China (31700582). We also thank Accdon (www.accdon.com) for their linguistic assistance during the preparation of this manuscript.

## Reference

Afsar U. Ahmed, Bryan R. G. Williams, Gregory E. Hannigan. 2015. Transcriptional Activation of Inflammatory Genes: Mechanistic Insight into Selectivity and Diversity. Biomolecules 5(4), 3087–3111.

Agnieszka Piatek, Zahir Ali, Hatoon Baazim, Lixin Li, Aala Abulfaraj, Sahar Al-Shareef, Mustapha Aouida and Magdy M. Mahfouz. 2015. RNA-guided transcriptional regulation in planta via synthetic dCas9-based transcription factors. Plant Biotechnology Journal 13, 578–589.

Bartsevich, V.V., Miller, J.C., Case, C.C. and Pabo, C.O. 2003. Engineered zinc finger proteins for controlling stem cell fate. Stem Cells 21, 632–637.

Bjorn Titz, Sindhu Thomas, Seesandra V. Rajagopala, et al. 2006. Transcriptional activators in yeast. Nucleic Acids Research 34(3), 955–967.

Blair, W.S., Bogerd, H.P., Madore, S.J. and Cullen, B.R. 1994. Mutational analysis of the transcription activation domain of RelA: identification of a highly synergistic minimal acidic activation module. Molecular and Cellular Biology 14, 7226–7234.

Blaise Weber, Johan Zicola, Rurika Oka, and Maike Stam. 2016. Plant Enhancers: A Call for Discovery. Trends in Plant Science 21(11), 974–987.

Douglas Vernimmen, Wendy A. Bickmore. 2015. The Hierarchy of Transcriptional Activation: From Enhancer to Promoter. Trends in Genetics 31, 696–708.

Huang T, Böhlenius H, Eriksson S, Parcy F, Nilsson O. 2005. The mRNA of the Arabidopsis gene FT moves from leaf to shoot apex and induces flowering. Science 309(5741), 1694–1696.

Kenneth S. Zaret, Jason S. Carroll. 2011. Pioneer transcription factors: establishing competence for gene expression. Genes & Development 25, 2227–2241.

Kooiker M., Xue GP. 2014. cDN A Library Preparation. In: Henry R., Furtado A. (eds) Cereal Genomics. Methods in Molecular Biology (Methods and Protocols), vol 1099. Humana Press, Totowa, NJ.

Kunzler, M., Braus, G.H., Georgiev, O., Seipel, K. and Schaffner, W. 1994. Functional differences between mammalian transcription activation domains at the yeast GAL1 promoter. EMBO Journal 13, 641–645.

Lakshmanan M, Kodama Y, Yoshizumi T, Sudesh K, Numata K. 2013. Rapid and efficient gene delivery into plant cells using designed peptide carriers. Biomacromolecules 14(1), 10–6.

Levi G. Lowder, Jianping Zhou, Yingxiao Zhang, Aimee Malzahn, Zhaohui Zhong Tzung-Fu Hsieh, Daniel F. Voytas, Yong Zhang, Yiping Qi. 2018. Robust Transcriptional Activation in Plants Using Multiplexed CRISPR-Act2.0 and mTALE-Act Systems. Molecular Plant 11, 245–256.

Li, L., Atef, A., Piatek, A., Ali, Z., Piatek, M., Aouida, M., Sharakuu, A., Mahjoub, A., Wang, G., Khan, S., Fedoroff, N.V., Zhu, J.K. and Mahfouz, M.M. 2013. Characterization and DNA-binding specificities of Ralstonia TAL-like effectors. Molecular Plant 6, 1318–1330.

Mahfouz, M.M., Li, L., Shamimuzzaman, M., Wibowo, A., Fang, X. and Zhu, J.K. 2011. De novo-engineered transcription activator-like effector (TALE) hybrid nuclease with novel DNA binding specificity creates double-strand breaks. Proc. Natl Acad. Sci. USA 108, 2623–2628.

Michaels SD, Amasino RM 1999. FLOWERING LOCUS C encodes a novel MADS domain protein that acts as a repressor of flowering. Plant Cell 11, 949–956.

Ohta, M., Matsui, K., Hiratsu, K., Shinshi, H. and Ohme-Takagi, M. 2001. Repression domains of class II ERF transcriptional repressors share an essential motif for active repression. Plant Cell 13, 1959–1968.

Ratcliffe OJ, Kumimoto RW, Wong BJ,Riechmann JL. 2003. Analysis of the *Arabidopsis* MADS AFFECTING FLOWERING gene family: MAF2 prevents vernalization by short periods of cold. Plant Cell 15, 1159–1169.

Remacle, J.E., Albrecht, G., Brys, R., Braus, G.H. and Huylebroeck, D. 1997. Three classes of mammalian transcription activation domain stimulate transcription in Schizosaccharomyces pombe. EMBO Journal 16, 5722–5729.

Rybakova KN, Bruggeman FJ,Tomaszewska A, Moné MJ, Carlberg C, Westerhoff HV. 2015. Multiplex Eukaryotic Transcription (In) activation: Timing, Bursting and Cycling of a Ratchet Clock Mechanism. PLoS Comput Biol 11(4): e1004236. doi:10.1371/journal.pcbi.1004236.

Scott JN, Kupinski AP, Boyes J. 2014. Targeted genome regulation and modification using transcription activator-like effectors. FEBS Journal 281(20), 4583–97.

Seipel, K., Georgiev, O. and Schaffner, W. 1992. Different activation domains stimulate transcription from remote (‘enhancer’) and proximal (‘promoter’) positions. EMBO Journal. 11, 4961–4968.

Sheldon CC, Burn JE, Perez PP, Metzger J, Edwards JA, Peacock WJ, Dennis ES. 1999. The FLF MADS box gene: A repressor of flowering in Arabidopsis regulated by vernalization and methylation. Plant Cell 11, 445–458.

Sumire Fujiwara, Shingo Sakamoto, Keiko Kigoshi, Kaoru Suzuki, Masaru Ohme-Takagi. 2014. VP16 fusion induces the multiple-knockout phenotype of redundant transcriptional repressors partly by Med25-independent mechanisms in Arabidopsis. FEBS Letters 588, 3665–3672.

Takahashi, K., and Yamanaka, S. 2006. Induction of pluripotent stemcells from mouse embryonic and adult fibroblast cultures by defined factors. Cell 126, 663–676.

Tian D, Xiang W, Chen X, et al. 2011. A long-term evaluation of biomass production in first and second rotations of Chinese fir plantations at the same site. Forestry: An International Journal of Forest Research 84(4), 411–418.

Tiwari, S.B., Belachew, A., Ma, S.F., Young, M., Ade, J., Shen, Y., Marion, C.M., Holtan, H.E., Bailey, A., Stone, J.K., Edwards, L., Wallace, A.D., Canales, R.D., Adam, L., Ratcliffe, O.J. and Repetti, P.P. 2012. The EDLL motif: a potent plant transcriptional activation domain from AP2/ERF transcription factors. Plant Journal 70, 855–865.

Triezenberg, S.J., Kingsbury, R.C. and McKnight, S.L. 1988. Functional dissection of VP16, the trans-activator of herpes simplex virus immediate early gene expression. Genes & Development 2, 718–729.

Wenjia Wang, Lianfeng Gu, Shanwen Ye, et al 2017. Genome-wide analysis and transcriptomic profiling of the auxin biosynthesis, transport and signaling family genes in moso bamboo (*Phyllostachys heterocycla*). BMC Genomics 18, 870.

Xian Xue, Qi Wang, Yanli Qu, Hongyang Wu, Fengqin Dong, Haoyan Cao, Hou-Ling Wang, Jianwei Xiao, Yingbai Shen and Yinglang Wan. 2017. Development of the photosynthetic apparatus of *Cunninghamia lanceolata* in light and darkness. New Phytologist 213, 300–313.

Xu Y, Liang Y, Yang M. 2019. Effects of Composite LED Light on Root Growth and Antioxidant Capacity of Cunninghamia lanceolata Tissue Culture Seedlings. Scientific Reports 9(1), 9766.

Yuan Y, Yang Y, Chen G. 2009. Fine root longevity of a Cunninghamia lanceolata plantation estimated by minirhizotrons. Journal of Subtropical Resources & Environment 4, 47–52.

Yueying Zhang, a,b Liang Du, a,1 Ran Xu, a Rongfeng Cui, a Jianjun Hao, c Caixia Sun, c and Yunhai Li. 2015. Transcription Factors SOD7/NGAL2 and DPA4/NGAL3 Act Redundantly to Regulate Seed Size by Directly Repressing KLU Expression in Arabidopsis thaliana. The Plant Cell 27, 620–632.

Zhihui Ma, Sizu Lin. 2019. Transcriptomic Revelation of Phenolic Compounds Involved in Aluminum Toxicity Responses in Roots of *Cunninghamia lanceolata (Lamb.)* Hook. Genes (Basel) 10(11), 835.

